# Polymorphism at four-fold degenerate site, but not at the intergenic regions, explains nucleotide compositional strand asymmetry in bacteria

**DOI:** 10.1101/2021.10.07.463495

**Authors:** Piyali Sen, Ruksana Aziz, Soumita Das, Nima Dondu Namsa, Ramesh Chandra Deka, Edward J Feil, Suvendra Kumar Ray, Siddhartha Sankar Satapathy

## Abstract

We investigated single nucleotide polymorphism in intergenic regions (IRs) and four-fold degenerate sites (FFS) in genomes of three γ-Proteobacteria and two Firmicutes to understand the mechanism of nucleotide compositional asymmetry between the leading and the lagging strands. Pattern of the polymorphism spectra were alike regarding transitions but variable regarding transversions in the IRs of these bacteria. Contrasting trends of complementary polymorphisms such as C→T vs G→A as well as A→G vs T→C in the IRs vindicated similar replication-associated strand asymmetry regarding cytosine and adenine deamination, respectively, across these bacteria. Surprisingly, the polymorphism pattern at FFS was different from that of the IRs and its frequency was always more than the IRs in these bacteria. Further, the polymorphism patterns within a bacterium were inconsistent across the five amino acids, which neither the replication nor the transcription-associated mutations could explain. However, the polymorphism at FFS coincided with amino acid specific codon usage bias in the five bacteria. Further, strand asymmetry in nucleotide composition could be explained by the polymorphism at FFS, not at the IRs. Therefore, polymorphisms at FFS might not be treated as nearly neutral unlike that in IRs in these bacteria.

## Introduction

As each of the four DNA bases can mutate to any of the three other bases, there are twelve possible directional substitution mutation types that include four transitions and eight transversions. These directional substitution mutations do not occur at equal frequencies in bacterial genomes for mechanistic reasons such as unequal stability among different base pairs, differential propensity of bases to damages such as deamination, oxidation, and radiation as well as selective reasons such as differential impact on the structure and function of DNA, RNA, and proteins. There are twice the number of transversions than transitions, but the observed frequency of transitions are double the transversions (Seplyarskiy et al. 2012; Duchêne et al. 2015; Lewis et al. 2016; Stoltzfus and Norris 2016; Lyons and Lauring 2017; Schroeder et al. 2017). Further, the higher propensity of deamination of C and oxidation of G bases increase the frequency of C→T and G→T base substitutions (Suzuki and Kamiya 2017), explaining the universal mutation bias towards A/T in genomes (Lind and Andersson 2008; Balbi et al. 2009; Hershberg and Petrov 2010; Hildebrand et al. 2010; Rocha and Feil 2010; Van Leuven and McCutcheon 2012).

Transition and transversion frequencies also vary between the leading strand (LeS) and the lagging strand (LaS) of replication known as strand substitution asymmetry (Lobry 1996; Frank and Lobry 1999), which is explained with GC (or AT) skew in chromosomes (Grigoriev 1998; Karlin et al. 1998; Rocha et al. 1999; Lobry and Sueoka 2002; Lobry 1996). Rapid deamination of cytosine to uracil in single stranded DNA leads to a higher rate of C→T mutation favouring a higher frequency of G and T in the LeS compared to the LaS (Rocha et al. 2006; Francino and Ochman 1997). The resulting GC (or AT) skew violates Chargaff’s second parity rule which states that the frequency of G and C (or A and T) should be approximately equal within individual DNA strands in genomes (Sueoka 1995; Forsdyke and Mortimer 2000; Powdel et al. 2009). Firmicutes are exceptional as they exhibit higher frequencies of A than T in the LeS (Rocha et al. 2006) implicating the selection and strong gene-orientation biases in the genomes of this bacterial group (Charneski et al. 2011). Assuming four-fold degenerate sites (FFS) evolve under near neutrality, Rocha et al. (2006) determined the polymorphism at four-fold degenerate sites (FFS) in seven different bacterial species to explain patterns of GC skew in genomes. They noted that relative consistency between taxa in terms of base compositional biases does not correspond with the underlying base substitution profiles. Although transcription-associated mutation was known to occur, the emphasis was given towards replication-associated mutation as highly expressed genes were not considered for analysis in their study (Davis 1989; Mugal et al. 2009; Kim and Jinks-Robertson 2012; Gaillard et al. 2013; Jinks-Robertson and Bhagwat 2014). Considering FFS to be nearly neutral, several researchers have extensively used FFS as a reference to describe strand as well as genome composition in bacteria (Muto and Osawa 1987; Rocha et al. 2006; Hershberg and Petrov 2010; Hildebrand et al. 2010), thereby neglecting the possible contribution of a selection mechanism. No cross-verification studies have been carried out in support of the above assumption of near neutrality of FFS being true. In case of nearly neutral nature, polymorphism patterns of different amino acids at FFS and at IRs are expected to be similar considering the role of context dependent mutation being minimal. The assumption of nearly neutral nature of FFS could be questioned based on the findings of earlier work in the areas of co-translational protein folding, selection on synonymous codons and ribosome mediated gene regulation (Satapathy et al. 2014; Sohmen et al. 2015; Ray and Goswami 2016; Ito 2016). Moreover, the findings by Charneski et al. (2011) support the role of selection for the atypical AT skew in *S. aureus*, emphasising the role of strand bias gene distribution in compositional skew (Charneski et al. 2011). A similar observation suggesting the selection at the IRs has been reported recently in bacteria (Thorpe et al. 2017). Considering mutation being AT biased in genomes (Hershberg and Petrov 2010; Hildebrand et al. 2010), it was previously assumed that the entire bacterial genome is under selection (Rocha and Feil 2010; Raghavan et al. 2012). The above findings have motivated us to relook at the polymorphism at FFS to determine the strand compositional asymmetry.

We have investigated nucleotide polymorphisms by analysing many strains of *Escherichia coli* (*Ec*)*, Klebsiella pneumoniae* (*Kp*), *Salmonella enterica* (*Se*) belonging to γ-Proteobacteria and two members of Firmicutes such as *Staphylococcus aureus* (*Sa*) and *Streptococcus pneumoniae* (*Sp*). The Firmicutes are known to exhibit different nucleotide compositional asymmetry between strands as compared to γ-Proteobacteria, which has been ascribed to replication-associated mutations due to the presence of two isoforms of DNA polymerase III alpha subunit, PolC and DnaE in Firmicutes (Rocha 2004; Saha et al. 2014). However, a large-scale analysis of diverse bacterial genomes indicates that purine asymmetry across two strands of replication, and different DNA polymerase compositions are neither essential nor exclusive features of the Firmicutes(Saha et al. 2014). It indicates a partial contribution of mutation in strand asymmetry composition in Firmicutes. Considering IRs are nearly neutral regions in a genome, polymorphism at IRs can be attributed to the replication-associated mutations (RAM) which are reflected by the contrasting pattern of complementary polymorphisms (e.g., C→T vs G→A; T→G vs A→C etc.) between the LeS and the LaS. Whereas the polymorphism at FFS can be attributed to RAM as well as transcription-associated mutations (TAM). In our comparative study, while the polymorphism at the IRs could be explained by RAM, polymorphism at FFS could not be explained by RAM or TAM in these bacteria, which was surprising. Instead, the polymorphism at FFS was observed to be influenced by the amino acid specific codon usage bias. Our work indicates that the polymorphism at FFS, not at IRs, can explain the nucleotide compositional strand asymmetry in bacterial genomes.

## Results

### Strand asymmetry in intergenic regions of bacteria due to transition polymorphisms

Prior to the analysis of nucleotide polymorphism at intergenic regions (IRs), we performed an elaborate study on its nucleotide composition. G+C% difference between whole genome and IRs was more prominent for the three γ-Proteobacteria (*Ec, Kp, Se*) than the two Firmicutes (*Sa*, *Sp*) (Supplementary Table 1). The G+C% in the IRs was similar between the LeS and the LaS within a bacterium. Abundance values between complementary nucleotides were more similar than that between two non-complementary nucleotides (Table 1). We found out different skews (AT/GC/KM/RY) in the IRs. AT skew was negative in the LeS and positive in the LaS in all bacteria except *Sa* where the reverse AT skew pattern was observed. The atypical AT skew in IR was in concordance with AT skew in *Sa* chromosome (Charneski et al. 2011). It is pertinent to note that the AT skew patterns were not consistent in IRs of *Sa* and *Sp*, although both belong to Firmicutes. In contrast to AT skew, GC skew values were found to be positive in the LeS and negative in the LaS of these bacteria. The magnitude of GC skew was observed to be higher than that of AT skew across the five bacteria. Among the five bacteria, the magnitude of both AT and GC skew was observed relatively high in *Sa*. The keto-amino (KM) and purine-pyrimidine (RY) skews were in general positive in the LeS and negative in the LaS of these bacteria (Table 1).

**Table 1:**
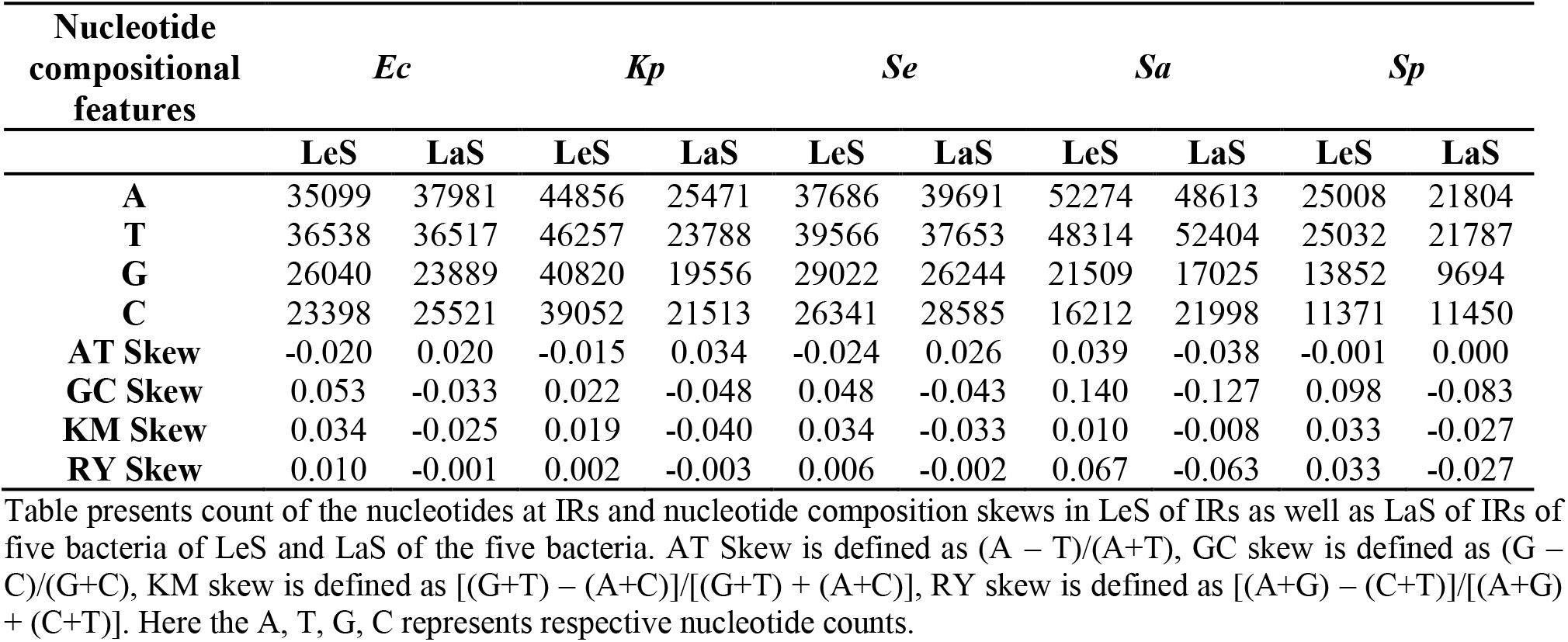
Compositional features of IRs in LeS and LaS in five bacteria.

The disparity of nucleotide composition between the LeS and the LaS was analysed in the context of nucleotide polymorphisms at the IRs. Frequencies of the twelve nucleotide polymorphisms were found out in the LeS and the LaS in these bacteria (Table 2). In general, transitions were more frequent than transversions. The *ti*/*tv* values varied from 1.5 to 2.3 among these bacteria (Table 2). In a more critical analysis of the transition and transversion frequencies, for example C→T vs. C→G in LeS, it was observed that the former was more than eight times than the latter in *Ec*, while the same was even ten time in *Se* and *Sp*. This fold differences were considerably higher than the expected value four-fold. The frequency values of complementary transition polymorphisms exhibited contrasting trends between the strands: C→T frequency was more than that of G→A in the LeS while the reverse was the case in the LaS; A→G frequency was more than that of T→C in LeS while the reverse was the case in the LaS (Figure 1). Frequency values of C→T and G→A were about two times more than that of A→G and T→C in both the strands (*p*-value < 0.01). This indicated that cytosine deamination is a major cause for the high frequency of C→T and G→A in genomes. Further, the higher frequency of C→T than that of G→A in LeS can be attributed to the higher propensity of single stranded DNA towards cytosine deamination over the double stranded DNA. Similarly, higher deamination of adenine in the single stranded DNA than the double stranded DNA might attribute towards the higher frequency of A→G than T→C in LeS. The difference between the frequencies of C→T and G→A within a strand was significantly more than that between A→G and T→C (*p*-value < 0.01). This finding is in concordance with both the following notions that cytosine is more prone to deamination than adenine in DNA and single stranded DNA has greater impact on cytosine deamination over adenine deamination. The strand asymmetry regarding transition polymorphisms observed in this study vindicates replication associated asymmetry in cytosine and adenine deamination between the LeS and LaS. As the pattern of transition polymorphism was similar across these bacteria, regarding the strand asymmetry, there seems to be similar impact of RAM in these two groups of bacteria.

**Table 2:** Polymorphism spectra at IRs.

| Substitution<br>spectra and<br>features | *Ec* |  | *Kp* |  | *Se* |  | *Sa* |  | *Sp* |  |
| --- | --- | --- | --- | --- | --- | --- | --- | --- | --- | --- |
|  | LeS | LaS | LeS | LaS | LeS | LaS | LeS | LaS | LeS | LaS |
| A→T | 0.023 | 0.024 | 0.022 | 0.023 | 0.024 | 0.026 | 0.036 | 0.035 | 0.014 | 0.014 |
| A→G | 0.053 | 0.045 | 0.048 | 0.041 | 0.085 | 0.079 | 0.048 | 0.037 | 0.049 | 0.044 |
| A→C | 0.015 | 0.017 | 0.014 | 0.016 | 0.02 | 0.027 | 0.012 | 0.014 | 0.015 | 0.016 |
| T→A | 0.024 | 0.024 | 0.022 | 0.023 | 0.022 | 0.026 | 0.038 | 0.033 | 0.016 | 0.014 |
| T→G | 0.018 | 0.014 | 0.015 | 0.013 | 0.024 | 0.023 | 0.016 | 0.011 | 0.015 | 0.017 |
| T→C | 0.045 | 0.054 | 0.042 | 0.048 | 0.071 | 0.095 | 0.042 | 0.044 | 0.046 | 0.05 |
| G→A | 0.091 | 0.114 | 0.093 | 0.105 | 0.175 | 0.228 | 0.122 | 0.134 | 0.126 | 0.181 |
| G→T | 0.036 | 0.033 | 0.04 | 0.036 | 0.057 | 0.065 | 0.051 | 0.051 | 0.043 | 0.051 |
| G→C | 0.012 | 0.015 | 0.017 | 0.017 | 0.018 | 0.021 | 0.016 | 0.017 | 0.013 | 0.015 |
| C→A | 0.035 | 0.037 | 0.037 | 0.039 | 0.057 | 0.067 | 0.058 | 0.045 | 0.046 | 0.045 |
| C→T | 0.112 | 0.098 | 0.095 | 0.088 | 0.218 | 0.188 | 0.15 | 0.115 | 0.187 | 0.127 |
| C→G | 0.014 | 0.013 | 0.015 | 0.015 | 0.02 | 0.02 | 0.02 | 0.014 | 0.014 | 0.015 |
| ti/tv | 1.701 | 1.757 | 1.527 | 1.549 | 2.269 | 2.145 | 1.466 | 1.500 | 2.318 | 2.150 |
| AT bias (ti) | 2.071 | 2.141 | 2.089 | 2.169 | 2.519 | 2.391 | 3.022 | 3.074 | 3.295 | 3.277 |
| AT bias (tv) | 2.152 | 2.258 | 2.655 | 2.586 | 2.591 | 2.640 | 3.893 | 3.840 | 2.967 | 2.909 |

Table represents 12 different substitution frequencies in LeS and LaS, for example C→T represents total C→T mutations divided by the total count of C in the gene. ti/tv is the ratio of transition to transversion. AT bias of transition is calculated as (C→T + G→A) / (A→G + T→C). AT bias of transversion is calculated as (G→T + C→A) / (T→G + A→C).

**Figure 1:**
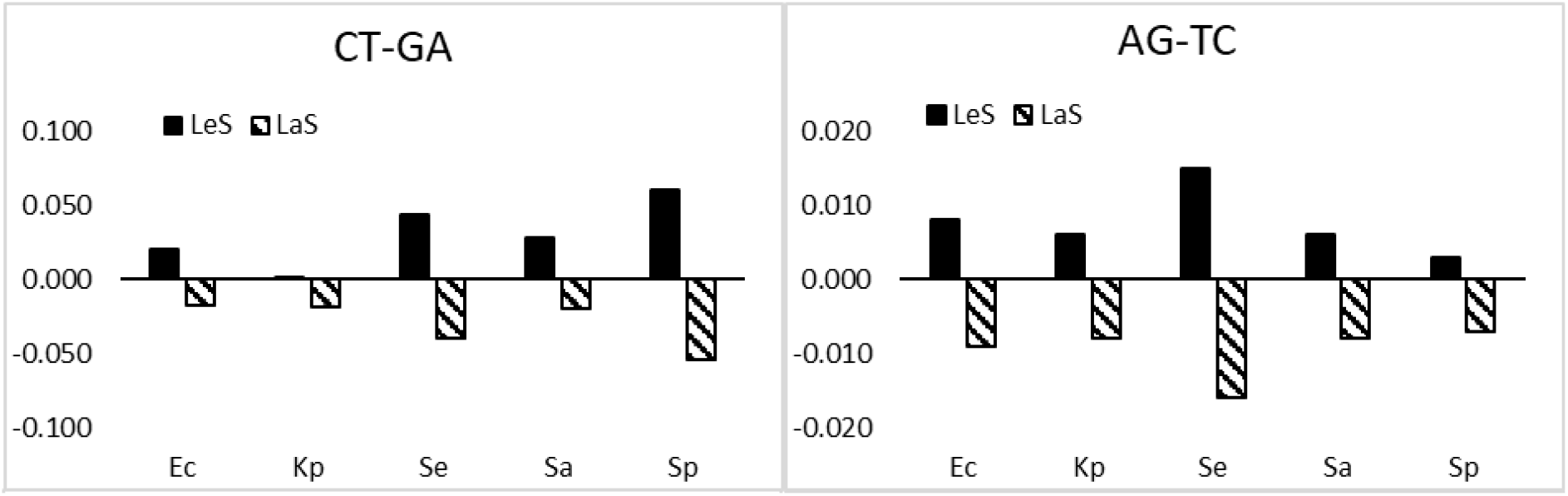
Difference between complementary transition polymorphisms in the LeS and the LaS at IRs. The figure represents a two-panel histogram of difference between complementary transition polymorphisms (C→T *vs* G→A and A→G *vs* T→C) in the LeS and LaS of IRs. The height of the vertical bar represents the polymorphism frequency difference between a complementary polymorphism pair. Black bar and striped bar represent polymorphism frequency differences in LeS and LaS, respectively. The X-axis represents the name of the five organisms: *Escherichia coli (Ec), Klebsiella pneumoniae (Kp), Salmonella enterica (Se), Staphylococcus aureus (Sa) and Streptococcus pneumoniae (Sp)*. Y-axis represents frequency difference values. In general, the pattern of complementary transition polymorphism in LeS and LaS are found to be in opposite direction.

Among the transversion polymorphisms, G→T (C→A) was the highest across these bacteria (Table 2). But, the order of the other polymorphisms regarding their frequencies was variable. For example, while A→T (T→A) frequency was the second highest among the transversions in *Ec*, *Kp*, *Se* and *Sa*; it was th lowest in *Sp*. Though, frequencies were different among *tv* polymorphisms, the frequency of a specific *tv* was similar between the strands. Therefore, the frequency values of complementary *tv* pairs were similar within a strand, unlike *ti*. The most frequent transversions G→T (C→A) is known to be due to the oxidation ofG to form 8oxoG (Loon et al. 2010), which seems to be occurring equally in both the strands. In *Sa* and *Sp* the frequency of G→T (C→A) was higher than the transitions polymorphisms A→G (T→C). This was contradicting the general notion that frequency of a transition polymorphism is higher than that of a transversion polymorphism. The reason behind the higher frequency of A→T (T→A) than that of A→C (T→G) or C→G (G→C) is not known. In *ti* as well as in case of *tv*, polymorphisms were more than two times biased towards A/T over G/C in three γ-Proteobacteria, and the same were more than three times biased towards A/T over G/C in two Firmicutes. This was in concordance with their genome composition values that polymorphism was more biased to AT in genome with low genome G+C composition.

In conclusion, polymorphism study at IRs has revealed that replication associated polymorphism asymmetry between the LeS and the LaS, is mainly attributed to deamination of cytosine and adenine. This polymorphism asymmetry is responsible for the positive GC or KM skew as well as the negative AT skew in the IRs of LeS and the *vice versa* in the LaS. From the frequency values of these polymorphisms, it could be found out that the polymorphism favours positive GC skew as well as negative AT skew in the LeS across these bacteria. The atypical AT-skew in *Sa*, could not be explained by the polymorphism spectra at IRs.

### The polymorphism pattern at the four-fold degenerate site (FFS) is different from that at IRs

Researchers had treated polymorphisms at FFS in bacteria as nearly neutral and had accordingly derived their conclusion regarding strand compositional asymmetry and genome composition (Muto and Osawa 1987; McLean et al. 1998; Reyes et al. 1998; Rocha and Danchin 2001; Rocha et al. 2006). However, there was no comparative study of polymorphism at FFS with that at IRs. Also, the polymorphisms were not compared across FFS of different amino acid codons within a bacterium. Here we found out polymorphism frequencies at FFS in codons of amino acids such as Val, Pro, Thr, Ala and Gly in the γ-Proteobacteria and Firmicutes (Supplementary Table 2). In general, *ti* was more frequent than *tv* across the five amino acids in these bacteria. However, the *ti*/*tv* values were variable, which indicated differences across these amino acid regarding polymorphism at FFS (Table 3). The transition as well as the transversion polymorphisms were more biased towards A/T in the Firmicutes than the γ-Proteobacteria (*p*-value < 0.01). Further the magnitudes of bias towards A/T over G/C were not consistent across the five amino acids either in the case of transition or in the case of transversion polymorphisms (Table 3). It is pertinent to note that polymorphisms such as C→T and G→A were not always more frequent than A→G and T→C. For example, in case of Thr in *Ec* and across the five amino acids in case of *Kp* (Supplementary Table 2). Further, among the transversion polymorphisms, G→T (C→A) were not the most frequent ones in *Ec* and *Kp*. So, the notion that C→T (G→A) is the most frequent transition polymorphism and G→T (C→A) is the most frequent transversion polymorphism is incorrect at FFS, unlike at IRs. Not only the frequency of a polymorphism was variable across the five amino acids, but the order of different polymorphisms regarding their frequency at FFS was also not consistent across the five amino acids (Supplementary Table 2, Figure 2). The difference among the amino acids regarding the polymorphisms within a bacterium indicated that the polymorphisms at FFS not necessarily be treated as nearly neutral.

**Table 3:** Comparison between transition-transversion polymorphism at FFS of five bacteria.

| Bacteria | Polymorphism<br>feature | Val |  | Pro |  | Thr |  | Ala |  | Gly |  |
| --- | --- | --- | --- | --- | --- | --- | --- | --- | --- | --- | --- |
|  |  | LeS | LaS | LeS | LaS | LeS | LaS | LeS | LaS | LeS | LaS |
| *Ec* | ti/tv | 1.558 | 1.589 | 1.288 | 1.342 | 1.598 | 1.739 | 1.513 | 1.521 | 1.732 | 1.893 |
|  | AT bias (ti) | 1.447 | 1.320 | 1.375 | 1.279 | 1.020 | 1.091 | 1.366 | 1.333 | 1.366 | 1.376 |
|  | AT bias (tv) | 0.915 | 0.850 | 1.117 | 0.945 | 0.758 | 0.795 | 0.904 | 0.921 | 1.027 | 1.096 |
| *Kp* | ti/tv | 1.701 | 1.761 | 1.283 | 1.464 | 1.707 | 1.755 | 1.631 | 1.647 | 1.733 | 1.787 |
|  | AT bias (ti) | 0.949 | 0.833 | 1.076 | 0.963 | 0.813 | 0.745 | 0.902 | 0.844 | 0.932 | 0.956 |
|  | AT bias (tv) | 0.944 | 0.866 | 0.900 | 0.946 | 0.973 | 0.766 | 0.927 | 0.846 | 0.777 | 0.905 |
| *Se* | ti/tv | 2.042 | 2.185 | 1.604 | 1.624 | 1.818 | 1.982 | 1.741 | 1.846 | 2.462 | 2.541 |
|  | AT bias (ti) | 2.292 | 2.165 | 2.202 | 2.242 | 1.843 | 2.037 | 2.195 | 2.098 | 1.559 | 1.508 |
|  | AT bias (tv) | 1.168 | 1.078 | 1.403 | 1.278 | 1.176 | 1.161 | 1.091 | 1.065 | 1.109 | 1.220 |
| *Sa* | ti/tv | 1.383 | 1.274 | 1.383 | 1.296 | 1.088 | 1.339 | 1.359 | 1.311 | 1.476 | 1.406 |
|  | AT bias (ti) | 3.362 | 2.922 | 5.130 | 5.917 | 4.041 | 4.786 | 3.928 | 4.491 | 3.077 | 2.713 |
|  | AT bias (tv) | 3.800 | 3.280 | 5.122 | 7.641 | 3.770 | 4.511 | 3.943 | 4.347 | 3.427 | 3.259 |
| *Sp* | ti/tv | 1.627 | 1.669 | 1.477 | 1.472 | 1.435 | 1.417 | 1.570 | 1.398 | 1.408 | 1.484 |
|  | AT bias (ti) | 2.097 | 2.225 | 3.967 | 4.051 | 2.663 | 2.852 | 2.619 | 2.817 | 2.698 | 3.168 |
|  | AT bias (tv) | 1.677 | 1.563 | 3.059 | 4.677 | 2.039 | 2.604 | 2.371 | 3.325 | 2.520 | 3.064 |

ti/tv is the ratio of transition to transversion. AT bias of transition is calculated as (C→T + G→A) / (A→G + T→C). AT bias of transversion is calculated as (G→T + C→A) / (T→G + A→C).

**Figure 2:**
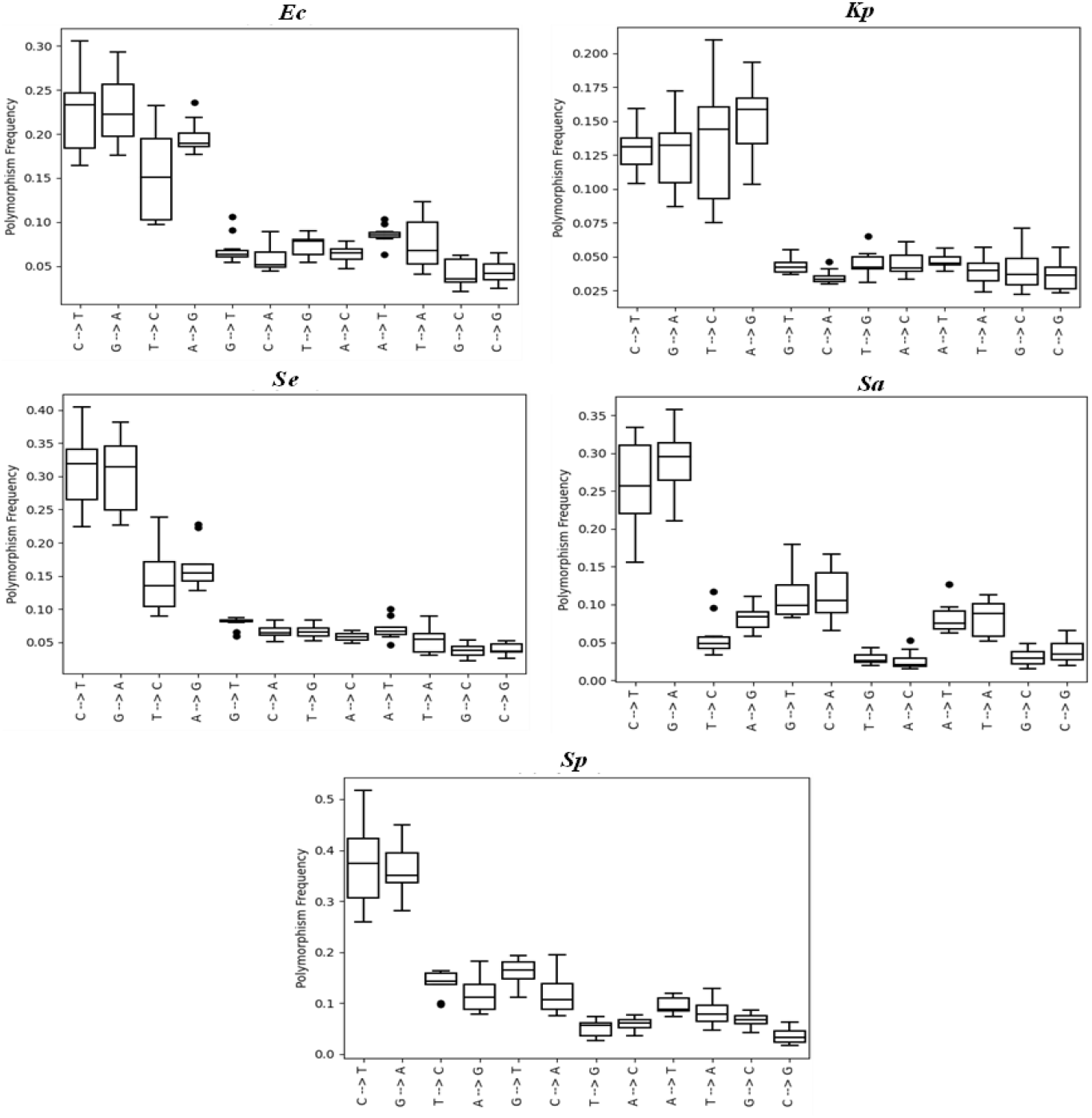
Polymorphism frequency at FFS across five amino acids in five bacteria. Figure presents distribution of 12 substitutions of FFS in terms of box-plot of Ec, Kp, Se, Sa and Sp. The x-axis presents the 12 substitutions and the y-axis presents the polymorphism frequency. The frequency of polymorphism is not uniform across the five amino acids.

We compared frequency values between complementary polymorphisms within a strand to find out any possible role of strand asymmetry. In general, the difference between C→T and G→A were positive in the LeS and negative in the LaS across the five amino acids in these five bacteria. This indicated replication associated strand asymmetry regarding cytosine deamination. Regarding the other transition polymorphisms, difference between A→G and T→C were high but non-uniform across the five amino acids within a bacterium: the difference can be positive in case of an amino acid while negative in case of another amino acid (Figure 3). Further, the difference between A→G and T→C values were similar both in the LeS and the LaS. This indicated that the difference was not generated due to replication associated strand asymmetry. Similarly, the inconsistent pattern across the amino acids indicated that the pattern was not due to transcription associated mutation. In case of complementary transversions, the difference values were high but not uniform across the five amino acids within a bacterium. Further, the difference values remain similar both in the LeS and the LaS. Unlike IRs, the difference between complementary polymorphisms were high at FFS and the difference value reflected were more of amino acid specific, not strand specific.

**Figure 3:**
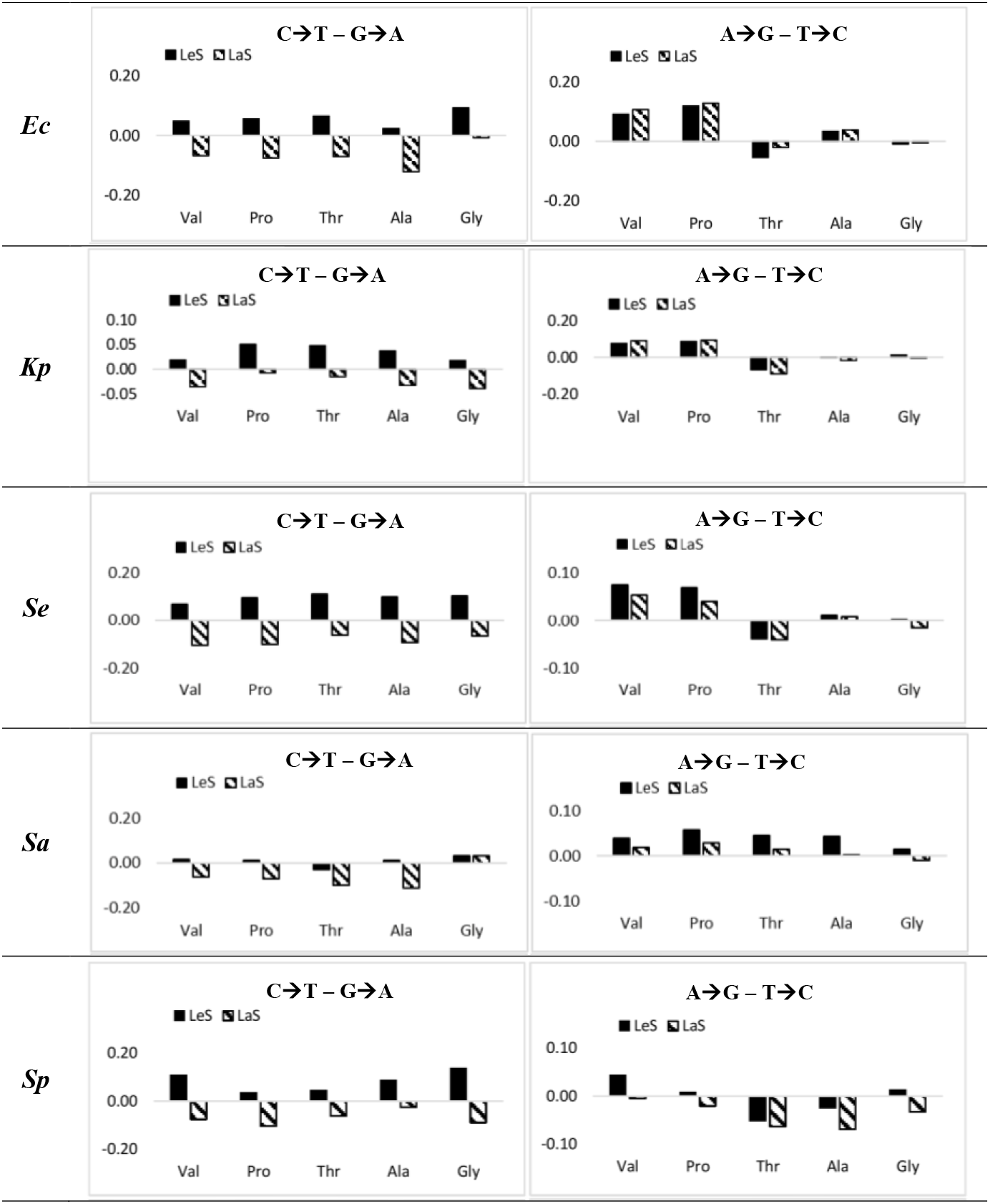
Difference between complementary transition polymorphisms in the LeS and the LaS at FFS. The figure presents a two-panel histogram of difference between complementary transition polymorphisms in the LeS and LaS at FFS of five bacteria. The height of the vertical bar represents the polymorphism frequency difference between a complementary polymorphism pair. Black bar and striped bar represent polymorphism frequency differences in LeS and LaS, respectively. The X-axis represents the name of the five amino acids namely Val, Pro, Thr, Ala and Gly. Y-axis represents frequency values. In general, the pattern of complementary transition polymorphism of C→T and G→A in LeS and LaS are found to be in opposite direction, whereas the other complementary transition polymorphism (A→G and T→C) do not show contrasting pattern.

### The polymorphism at the four-fold degenerate site coincides with codon usage bias

The amino acid specific polymorphisms at FFS indicated that the polymorphisms were most probably influenced by codon usage bias. So, we compared polymorphisms with nucleotide frequency at FFS of the five amino acids, which represented the codon usage bias of individual amino acids. Wide variation regarding nucleotide frequency at FFS was observed across the five amino acids within a genome (Table 4). In general, codon usage bias was observed to be strand independent as the frequency values were similar between the two strands for an amino acid at FFS. This was in concordance with observations by earlier researchers (Sharp et al. 2005; Shah and Gilchrist 2011; Wald et al. 2012). However, a moderate impact of strand asymmetry was observed on codon usage bias because in general the frequencies of G and T at the FFS of an amino acid in LeS was more than that in LaS. The reverse was true regarding the frequencies of A and C.

**Table 4:**
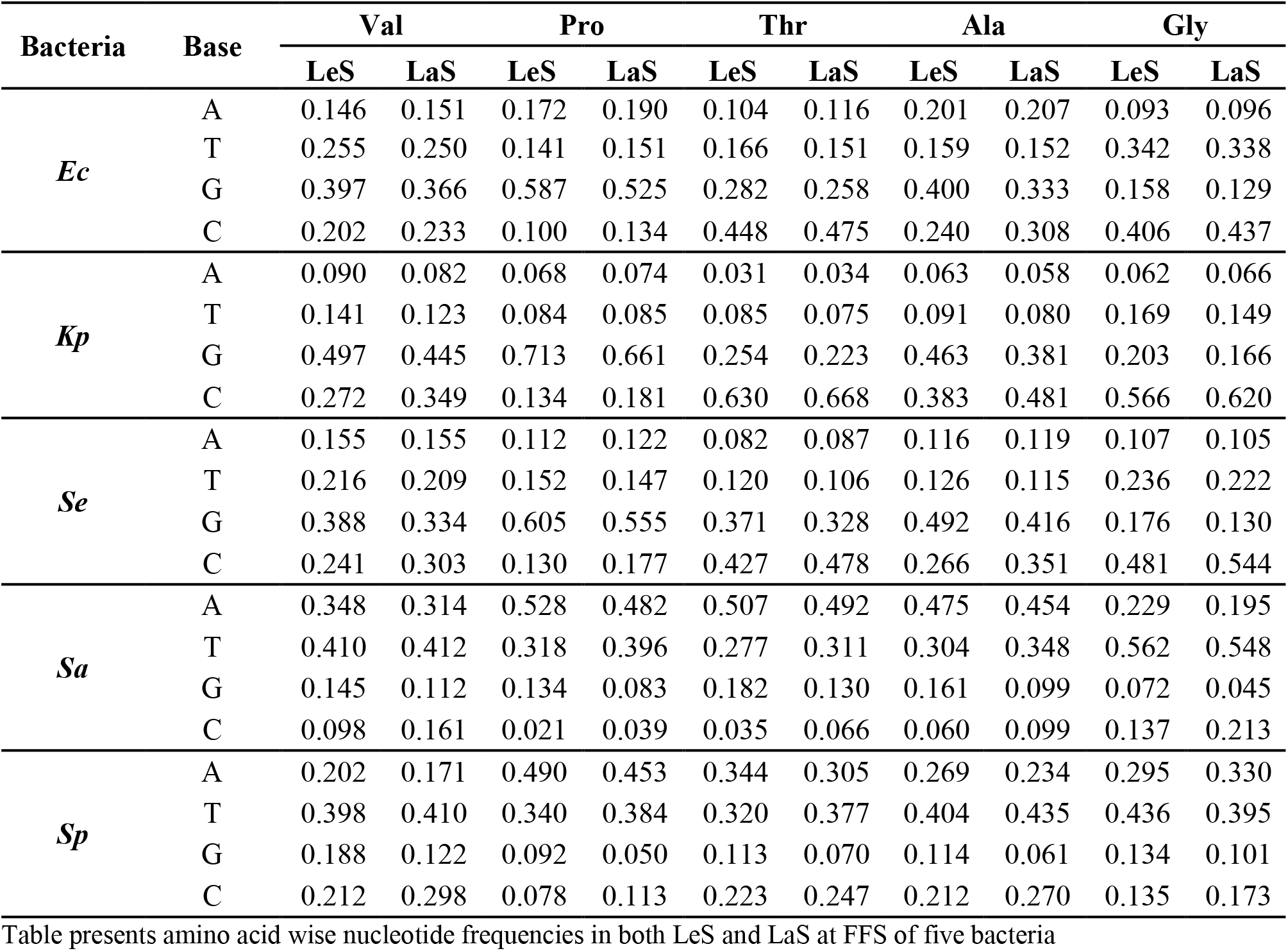
Nucleotide frequency at FFS.

| Bacteria | Base | Val |  | Pro |  | Thr |  | Ala |  | Gly |  |
| --- | --- | --- | --- | --- | --- | --- | --- | --- | --- | --- | --- |
|  |  | LeS | LaS | LeS | LaS | LeS | LaS | LeS | LaS | LeS | LaS |
| *Ec* | A | 0.146 | 0.151 | 0.172 | 0.190 | 0.104 | 0.116 | 0.201 | 0.207 | 0.093 | 0.096 |
|  | T | 0.255 | 0.250 | 0.141 | 0.151 | 0.166 | 0.151 | 0.159 | 0.152 | 0.342 | 0.338 |
|  | G | 0.397 | 0.366 | 0.587 | 0.525 | 0.282 | 0.258 | 0.400 | 0.333 | 0.158 | 0.129 |
|  | C | 0.202 | 0.233 | 0.100 | 0.134 | 0.448 | 0.475 | 0.240 | 0.308 | 0.406 | 0.437 |
| *Kp* | A | 0.090 | 0.082 | 0.068 | 0.074 | 0.031 | 0.034 | 0.063 | 0.058 | 0.062 | 0.066 |
|  | T | 0.141 | 0.123 | 0.084 | 0.085 | 0.085 | 0.075 | 0.091 | 0.080 | 0.169 | 0.149 |
|  | G | 0.497 | 0.445 | 0.713 | 0.661 | 0.254 | 0.223 | 0.463 | 0.381 | 0.203 | 0.166 |
|  | C | 0.272 | 0.349 | 0.134 | 0.181 | 0.630 | 0.668 | 0.383 | 0.481 | 0.566 | 0.620 |
| *Se* | A | 0.155 | 0.155 | 0.112 | 0.122 | 0.082 | 0.087 | 0.116 | 0.119 | 0.107 | 0.105 |
|  | T | 0.216 | 0.209 | 0.152 | 0.147 | 0.120 | 0.106 | 0.126 | 0.115 | 0.236 | 0.222 |
|  | G | 0.388 | 0.334 | 0.605 | 0.555 | 0.371 | 0.328 | 0.492 | 0.416 | 0.176 | 0.130 |
|  | C | 0.241 | 0.303 | 0.130 | 0.177 | 0.427 | 0.478 | 0.266 | 0.351 | 0.481 | 0.544 |
| *Sa* | A | 0.348 | 0.314 | 0.528 | 0.482 | 0.507 | 0.492 | 0.475 | 0.454 | 0.229 | 0.195 |
|  | T | 0.410 | 0.412 | 0.318 | 0.396 | 0.277 | 0.311 | 0.304 | 0.348 | 0.562 | 0.548 |
|  | G | 0.145 | 0.112 | 0.134 | 0.083 | 0.182 | 0.130 | 0.161 | 0.099 | 0.072 | 0.045 |
|  | C | 0.098 | 0.161 | 0.021 | 0.039 | 0.035 | 0.066 | 0.060 | 0.099 | 0.137 | 0.213 |
| *Sp* | A | 0.202 | 0.171 | 0.490 | 0.453 | 0.344 | 0.305 | 0.269 | 0.234 | 0.295 | 0.330 |
|  | T | 0.398 | 0.410 | 0.340 | 0.384 | 0.320 | 0.377 | 0.404 | 0.435 | 0.436 | 0.395 |
|  | G | 0.188 | 0.122 | 0.092 | 0.050 | 0.113 | 0.070 | 0.114 | 0.061 | 0.134 | 0.101 |
|  | C | 0.212 | 0.298 | 0.078 | 0.113 | 0.223 | 0.247 | 0.212 | 0.270 | 0.135 | 0.173 |

Table presents amino acid wise nucleotide frequencies in both LeS and LaS at FFS of five bacteria

To understand the relation of the polymorphism with codon usage bias, if any, we did a detailed comparative study in individual bacteria. The polymorphism spectra at FFS for individual bacterium is given in Supplementary Table 2. In case of *Ec*, the polymorphism pattern between complementary nucleotide pairs revealed that Pro and Gly often behaved opposite to each other. Considering the previous knowledge that G-ending codon in Pro (CCG) is the most preferred whereas the G-ending codon (GGG) in Gly is the least preferred (Satapathy et al. 2016), we analysed both forward and reverse nucleotide polymorphisms. In case of Gly, G→C and G→T transversions were more frequent than C→G and T→G respectively, that supported the higher abundance of GGT/GGC over GGG in the genome. Whereas in case of Pro, G→C and G→T transversions were found to be less frequent than C→G and T→G respectively, that supported the lower abundance of CCT/CCC than CCG in the genome. A→T was more frequent than T→A in Val and Gly that favoured higher abundance of GTT/GGT over GTA/GGA. A→T was less frequent than T→A in Pro that favoured CCA to be more frequent than CCT. T→C was the most frequent in case of Thr among the five amino acids that made ACC the most preferred codon. Low frequency of T→C in case of Pro and Val, favoured the low frequency of GTC/CCC. While C→T and T→C were of similar frequency in case of Thr, C→T was more than the frequency of T→C in case of Val, Pro and Ala that favoured the higher abundance of T-ending codon (GTT/CCT/GCT) over the C-ending codons. G→A transition was more frequent than A→G in case of Gly, which corresponds to the higher frequency of GGG over GGA. These observations suggested that polymorphism at FFS was influenced by codon usage bas in *Ec*.

A similar comparative study of the polymorphism at FFS and codon usage bias in these amino acids were studied in the other two γ-proteobacteria *Kp* and *Se*. In *Kp*, as the genome is G+C high, G/C-ending codons are more abundant over the A/T-ending codons. In general A→G was more frequent than C→T and G→A in all amino acids that favoured higher abundance of G-ending codons in the genome. A→C was more frequent than C→A in all amino acids except Pro that supported CCC being preferred low here. Further, we noticed correlation in preference of C-ending codons and higher frequency of G→C in Thr and Gly, whereas the G-ending codons were preferred in Val and Pro that corresponded to the higher frequency of C→G. A→T was more frequent than T→A in Val and Gly, which supported the T-ending codons being preferred over the A-ending codons in these amino acids. T→G was more frequent than G→T in Pro while T→G was less frequent than G→T in Gly which supported G-ending codon being preferred in Pro but not in Gly. Therefore, the impact of codon usage bias was observed on the polymorphism at FFS of *Kp*. Similarly, in *Se*, G→C transversion was more frequent than C→G in Thr and Gly while the reverse pattern was true in Pro in both the strands. This was in concordance with the preference of C-ending codon in Thr, avoidance of C-ending codon in Pro and G-ending codon in Gly. Polymorphism at A→T was more frequent than T→A in Val and Gly which supported the higher abundance of GTT/GGT in the genome. Therefore, polymorphism pattern at FFS was influenced by codon usage bias in *Se*.

In the two Firmicutes, A- and T-ending codons were the most frequent codons across the five amino acids. In *Sa*, A-ending codons were more frequent than T-ending codons in Pro, Thr and Ala. In concordance, A→T was more frequent than T→A in Val and Gly, while the reverse was the case in Pro, Thr and Ala. In addition, G→A was more frequent than C→T in case of Thr because ACA is the most frequent codon here. In *Sp*, A-ending codon was more frequent than T-ending codons in Pro. It was obvious to observe that A→T was more frequent than T→A in case of Val, Ala and Gly but the reverse was true in case of Pro, which was in concordance with the codon usage bias. Therefore, polymorphism at FFS in *Sa* as well as in *Sp* was influenced by codon usage bias in these bacteria.

## Discussion

Analysing nucleotide polymorphism in genome sequences of several strains belonging to a single species provides avenue to study mechanisms of molecular evolution. Further the leading and the lagging strands in chromosomes can be easily segregated using computational method facilitating researcher to investigate the replication-associated mutation asymmetry between the strands in bacteria (Frank and Lobry 1999). Despite of low abundance of IRs, these regions can be distinctly identified because of the simplicity of the coding sequences in bacterial chromosomes. In this study, nucleotide polymorphism has been analysed at the intergenic regions (IRs) and four-fold degenerate site (FFS) of three bacterial species namely *E. coli* (*Ec*), *K. pneumoniae* (*Kp*)*, S. enterica* (*Se*) belonging to γ-proteobacteria and two other bacterial species namely *S. aureus* (*Sa*) and *S. pneumoniae* (*Sp*) belonging to Firmicutes. Nucleotide compositional asymmetry between the strands is well studied in bacterial chromosomes (Lobry 1996; Lobry and Sueoka 2002). The higher abundance of the keto nucleotides (G, T) in the LeS than the LaS has been explained based on frequent cytosine deamination in single stranded DNA (Reyes et al. 1998; Frank and Lobry 1999; Rocha 2004) which is corroborated by the observation of positive GC skew and negative AT skew in the LeS in bacteria. However, the higher magnitude of GC skew than AT skew in the LeS of most bacterial genomes could not be explained based on the cytosine deamination theory. Rocha et al. (2006) had explained different mutation bias that might lead to similar skew patterns in bacterial genomes. Similarly, the positive AT skew observed in the LeS of *S. aureus* could not be explained by the cytosine deamination theory. Therefore, a detailed investigation has been done in this study to understand the nucleotide compositional asymmetry between the strands in bacteria. Polymorphism analysis at IRs of these five bacteria has revealed that C→T and G→A transition polymorphism are the most frequent ones and display the main difference between the strands. This is in favour of the notion that cytosine deamination is the major cause of polymorphism in genomes and the process is more frequent in the LeS than the LaS. In a small magnitude, we have also observed that A→G and T→C contributes towards strands asymmetry at IRs. In parallel with cytosine deamination theory, it may be hypothesized that more frequent adenine deamination in LeS might result in higher A→G transition in the strand than the complementary strand. It is known that cytosine deamination and adenine deamination have an opposite impact on genome G+C%. Regarding transition polymorphisms at IRs, the five bacteria behave similar in this study. However, in transversions, frequency of a polymorphism is observed to be similar between the strands and the difference value between complementary transversions within strand is very low. Therefore, contribution of transversion polymorphism in strand asymmetry is very low in the IRs of these bacteria. It is pertinent to note that frequency of G→T (C→A) are higher than other transversions. Transversion polymorphisms increases A/T at IRs like transition polymorphisms. But the bias towards A/T of these polymorphisms are more in the two Firmicutes than the γ-Proteobacteria. The G→T (C→A) value is higher as well as A→G (T→C) value is lower in Firmicutes in comparison to the γ-Proteobacteria for which A/T bias is observed to be more in the former than the latter. The polymorphism study at the IRs suggests that, the replication associated strand asymmetry is indifferent between the two groups of bacteria. Therefore, the atypical AT skew in the chromosome of *Sa* is not supported by the polymorphism at IRs.

In the two Firmicutes, *Sa* and *Sp* exhibit opposite patterns of nucleotide composition at FFS. The nucleotide A is more frequent than T in *Sa*, while T is more frequent than A in *Sp*. The coding sequence is more abundant in the leading strand than the lagging strand of the Firmicutes (Rocha 2004). Therefore, the abundance of A is more than T in the LeS of *Sa* and the abundance of T is more than A in the LeS of *Sp*. Codon usage bias at FFS of an amino acid is similar between the strands indicating the weak influence of strand specific polymorphism. Therefore, the atypical AT skew in *Sa* can be attributed to codon usage bias, which is due to the selection on codon usage bias. Our findings are in concordance with the earlier observation that selection and gene distribution asymmetry between the strands was associated with the atypical AT skew in *Sa* (Charneski et al. 2011). Hence, our observations in this study suggest that selection on codon usage bias influences the polymorphism at FFS in the Firmicutes. In conclusion, we would like to state that, our understanding regarding the influence of codon usage bias on the compositional strand asymmetry become clear in this study because of the polymorphism study done separately in IRs and FFS.

Genome G+C% in bacteria varies from 13.0 to 75.0% which accounts a difference of 62% between the minimum and maximum genome composition (Raghavan et al. 2012). Directional mutation bias in support of the neutral theory of evolution has been proposed to explain genome G+C% in organisms (Sueoka 1988). In support of directional mutation theory, Muto and Osawa (1987) demonstrated that synonymous codon usage varies between high and low G+C bacterial genomes. But theoretical analysis of the G+C% of the synonymous codons suggests that the maximum G+C composition difference between two synonymous codons (e.g., GGU and GGC) of an amino acid can be 33.33% with exceptions only in Arg (e.g., CGG, AGU) and Leu (e.g., UUA, CUG) where the difference can be up to 66.67%. Hence, the synonymous codon usage range should be held accountable to a value around 33.33% instead of 62.0% (75.0 – 13.0). It can be argued that the directional mutation theory inadequately explains the genome G+C composition in bacteria as IRs contribute a minor portion of the genome size. It is pertinent to note that the results of earlier mutation analysis in bacterial genomes could not provide substantial evidence in support of the directional mutation theory (Hershberg and Petrov 2010; Hildebrand et al. 2010; Rocha and Feil 2010). Hence, the possible existence of an unknown selection mechanism responsible for genome G+C% has been hypothesized (Rocha and Feil 2010; Raghavan et al. 2012). The role of recombination in determining the genome composition of bacteria have also been implicated (Bobay and Ochman 2017). G+C% variation at FFS across the amino acids in different genomes reported in this work is an indication of codon usage bias influencing the observed difference in genome composition. Notably, we found that Pro, Thr and Ala codons having C at the second position differ from each other in terms of codon usage bias. Similarly, Val, Ala and Gly codons having G at the first position also differ from each other in terms of codon usage bias. Therefore, the possibility of context-dependent polymorphism causing codon usage difference across the amino acids is not anticipated. The polymorphism difference observed across the amino acids at FFS suggest that the difference is due to the amino acid specific translational selection. Our comparative analysis of polymorphism and codon usage bias in the five studied bacteria led us to believe that the selection on codon usage bias responsible for the observed polymorphism. It is already known that GGG and CCC codons are prone to translational frameshift (O’Connor 1998, 2002). Interestingly, GGG and CCC codons were not preferred in both the strands of five studied bacteria. Transversions such as G→C is more than C→G in case of Gly while the same is less in case of Pro. GGC codon has been reported to be selected positively in bacterial genomes (Satapathy et al. 2014, 2016). Further amino acid specific codon usage bias is similar in the two strands indicating a weak influence of the strand specific mutation bias, in comparison to the translational selection in genomes. This observation supports that G+C% variations across different amino acids at FFS is due to selection on codon usage bias and may not be due to the directional mutation. Further, considering a limited set of high expression genes, we observed that the polymorphism at FFS of the high expression genes is in line with that at FFS of whole genome. It is interesting to note that the earlier notion of genome composition determining the codon usage bias (Muto and Osawa 1987) is found inconsistent in this study. Now our analysis using large number of genomes of γ-Proteobacteria and Firmicutes have suggested the role of codon usage bias in determining the genome G+C% supporting the selection theory of evolution. Assuming that the selection theory is true, it is speculated that in an AT rich genome, A/T ending synonymous codons are likely to be selected over the G/C ending synonymous codons and *vice versa* is true for GC rich genomes (Hershberg and Petrov 2009). We anticipated that future research on translation rate of synonymous codons is expected to uncover the mechanism of genome composition in AT and GC rich bacteria.

Large range of genome G+C% is a classic example of the neutral theory of evolution in bacteria which means that there is no specific advantage that could be linked to genome composition (Lassalle et al. 2015). Under this assumption, the low genomic G+C content of endosymbiotic bacteria were considered in favour of the neutral theory. However, recently Dietel et al. (2019) have discussed that selective advantages favour high genomic AT-contents in intracellular genetic elements. Genome composition is known to be associated with bacterial phylogeny such as Firmicutes with low genome G+C%, Actinomycetes and β-Proteobacteria with high genome G+C% (Satapathy et al. 2010). However, the reason for the high and low G+C% in these phyla is not clearly understood. But phylogeny specific optimal codon selection has been reported recently (Satapathy et al. 2016). Future understanding of translational decoding by the ribosome might explain the phylogeny specific codon usage bias and genome composition. It is pertinent to note that ribosome mediated gene regulation by co-translational protein folding has been demonstrated to be species specific in *E. coli* and *B. subtilis* (Sohmen et al. 2015). It has been reported that genome size and G+C% are positively correlated (Satapathy et al. 2010). However, we have observed a lack of correlation between the genome size and genome G+C% in different phylogeny (Supplementary Table 3). It has been reported that the strength of selection on codon usage bias is variable among the bacteria (Sharp et al. 2005; Satapathy et al. 2012, 2014, 2016). In that case bacteria with poor selection on codon usage bias should exhibit low genome G+C%, while bacteria with high genome G+C% must exhibit high selection on codon usage bias. In a different study, it has been shown that bacteria with high genome G+C% indeed exhibited selection on codon usage bias (Satapathy et al. 2014). Hence, this further supports that the selection of codon usage bias is responsible for genome composition in bacteria. As codon usage bias is universal in bacteria, it may be possible that the difference between two genomes regarding codon usage bias may act as a selection against lateral gene transfer in bacteria.

## Materials and Methods

### Segregating the leading and the lagging strands in bacterial chromosomes

We carried out a detailed single nucleotide polymorphism study using computational analysis of genomes of total 157 *Escherichia coli* (*Ec*) strains (Thorpe et al. 2017), 208 *Klebsiella pneumoniae* (*Kp*) strains (Holt et al. 2015), 366 *Salmonella enterica* (*Se*) strains (Thorpe et al. 2017), 132 *Staphylococcus aureus* (*Sa*) strains (Reuter et al. 2016) and 264 *Streptococcus pneumoniae* (*Sp*) strains (Chewapreecha et al. 2014). Considering cumulative GC skew diagram for each bacterium, we segregated respective chromosomes into the leading strand (LeS) and the lagging strand (LaS) as has been described earlier by different researchers (Lobry 1996; Grigoriev 1998). The method is mentioned below in brief. We found out abundance values of each nucleotide along a genome sequence using non-overlapping moving window of size 1.0 kb. GC skew was calculated as (G-C)/(G+C). Similarly AT skew was calculated as (A-T)/(A+T), RY skew was calculated as [{(A+G)-(C+T)}/(A+G+C+T)], and KM skew was calculated as [{(G+T)-(A+C)}/(A+G+C+T)]. For each of the bacteria, cumulative skew diagrams were generated from these deviations, which was used to identify the leading (LeS) and the lagging (LaS) strands regions in a chromosome (Supplementary Figure 1a and 1b). GC skew is positive in the LeS whereas the same is negative in the LaS. Similarly, KM skew is positive in the LeS whereas the same is negative in the LaS. A schematic view of the LeS and the LaS in a double stranded DNA is presented in Figure 4. The LeS and the LaS regions of the Watson strand are aligned with the LaS and the LeS, respectively, of the Crick strand in chromosomes. Using the coordinates of protein coding genes (CDS) from the genome annotation, the CDS were mapped to the LeS and the LaS. Sequences other than those coding for the rRNA, tRNA, protein genes and misc_RNAs were considered as intergenic regions (IRs). IRs in the LeS and the LaS of the Watson strand are aligned opposite to IRs in the LaS and the LeS, respectively, of the

**Figure 4:**
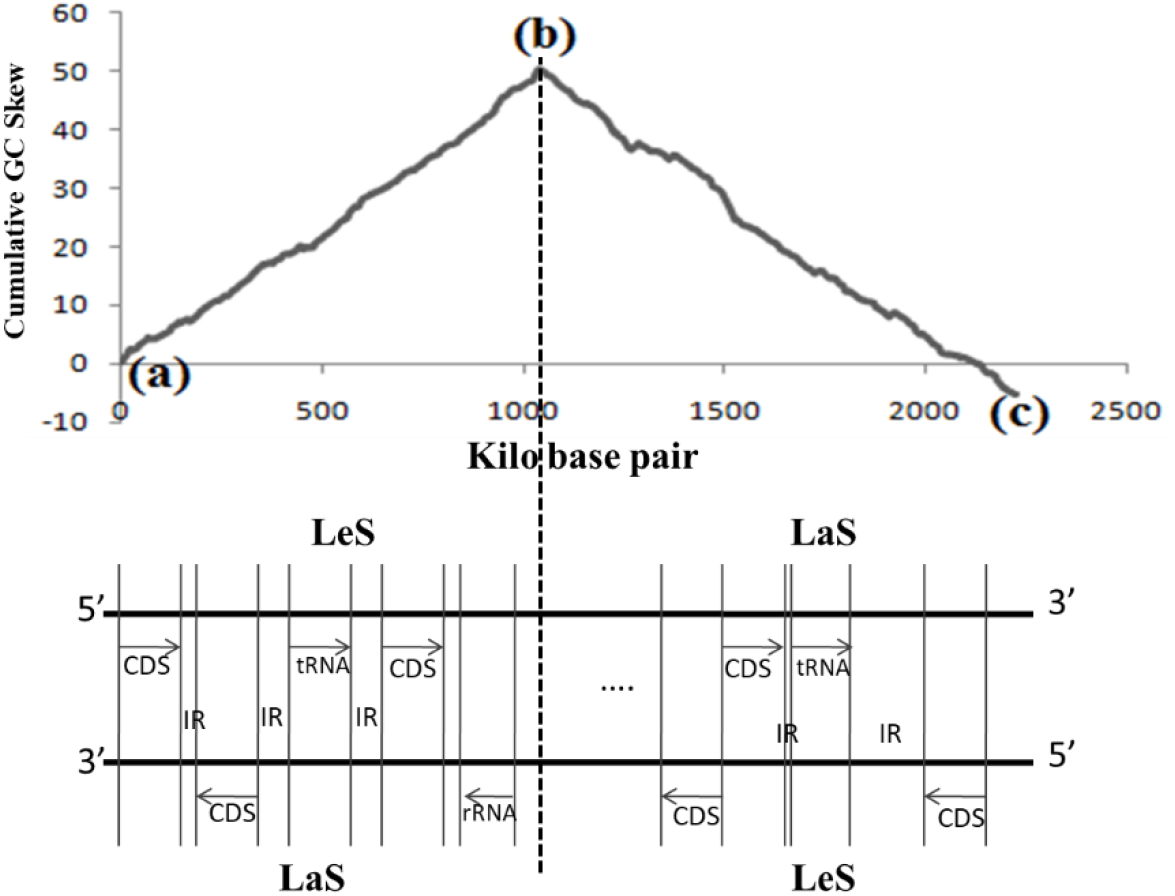
A schematic view of the distribution of IRs, CDS, tRNA and rRNA in the leading and lagging strands in double stranded DNA. Figure represents a schematic view of the LeS/LaS and distribution of IRs and CDS, tRNA and rRNA in a double stranded DNA. Considering nucleotide composition of the Watson strand, the skew diagram is generated. In the strand, the region between point (a) and point (b) with positive GC skew is designated as the LeS and the sequence between point (b) and point (c) as the LaS. The LeS and the LaS regions of the Watson strand are aligned with the LaS and the LeS, respectively, of the Crick strand in chromosomes.

Crick strand (Figure 4). For polymorphism analysis, IRs in either the Watson or the Crick strand were considered. IRs belonging to either the Watson or the Crick strand were segregated into the LeS and the LaS (Supplementary Figure 1a, 1b and Figure 4). In our analysis, we included only large IRs (size greater than 100 bases) from where 35 bases from both the ends of each IR were ignored to minimize the inclusion of any regulatory regions and considered only the nearly neutral polymorphisms.

### Polymorphism analysis at IRs and at FFS in CDS of the bacterial chromosomes

For each bacterium, we extracted alignments of intergenic regions (IRs) and protein-coding regions (CDS) using computer programs written in Python script. Considering the most frequent nucleotide at a position in the sequence alignment, we computed a consensus sequence which was used to identify polymorphisms at different positions. A detailed description of the approach used for analysing polymorphisms in this study is provided in the Supplementary Document 1. Phylogenetic relationships among the five bacteria (Supplementary Figure 2a) were obtained using the reference sequence of *rpoB* and *rpoC* genes by the MEGAX software (Kumar et al. 2018). The three γ-proteobacteria (*Ec*, *Kp* and *Se*) and two Firmicutes (*Sa* and *Sp*) made different clusters. Further, phylogenetic relationships among the population (Supplementary Figure 2b-2f) were constructed using the *rpoB* sequence of all strains of individual organisms (Kumar et al. 2018). The distribution of polymorphisms among the strains in reference to *rpoB* and *rpoC* genes (Supplementary Figure 3) shows that, in general, polymorphism observed in this study is not because of any specific strain but mutations accumulated among all the strains (Supplementary Table 4). A known set of previously published high expression genes (Sharp et al. 2005; Sen et al. 2020) were considered in this work for the analysis of nucleotide composition and polymorphism at FFS in high expression genes (HEGs) (Supplementary Table 5 and 6).

In coding sequences (CDS), we considered polymorphism at FFS of the amino acids such as Val, Pro, Thr, Ala and Gly. For example, if a nucleotide change (suppose A→T) observed at the 3^rd^ position of codon, the corresponding codon was found out in the reference sequence (considering the preceding two nucleotides). If the codon codes for Val (i.e., the codon is GTT/GTC/GTA/GTG), then we increase A→T change for Val by 1. Using this approach, we calculated polymorphism at FFS of the five amino acids. Polymorphism frequencies were normalized by dividing the total count of a given change by the total count of the nucleotide in which polymorphism has occurred in the reference sequence. For example, if the total number of C→T change is 20 and the total number of C in the reference sequence (either at IRs or at FFS of that amino acid) is 100, then the normalized frequency is calculated as 20/100 = 0.2. The frequencies of different nucleotide polymorphisms were calculated accordingly. For statistical analysis and determining p-value for significance test, Mann Whitney test is used (Mann and Whitney 1947).

## Acknowledgements

We are grateful to Dr. Harry Thorpe for his kind input and useful discussion on the genome sequence data. PS is thankful to the University Grants Commission, GoI, New Delhi (Award letter No. 3314/(NET-JULY 2016) for the fellowship. RA is thankful to Tezpur University and DBT, GoI (Grant No. BT/511/NE/TBP/2013) for the fellowship. SKR, SSS and RCD thank the DBT, GoI, New Delhi for the project grants in the area computational biological research (BT/511/NE/TBP/2013, BT/PR16361/NER/95/192/2015, BT/PR16182/NER/95/92/2015). SSS is thankful to DBT for the North East Overseas Associateship that helped him to work at the University of Bath. The authors are thankful to SMBE for holding the satellite meeting at Kaziranga, Assam, India on Dec 14^th^-16^th^, 2017. SSS, RCD, SKR and NDN duly acknowledged the support of DBT funded Centre for Bioinformatics at Tezpur University. We thank Dr. S. Chatterjee, the organizer of M3II 2019, India where we received useful comments on this work. We also thank Dr. Jagreet Kaur, Dr. P.B Patil and Prof. S. K. Kar for discussion on the manuscript.

